# Batch Effect Correction in a Functional Colorectal Cancer Organoid Clinical Correlation Study

**DOI:** 10.64898/2026.02.05.704065

**Authors:** Gavin R. Oliver, António Miguel de Jesus Domingues, Carlton C. Barnett

## Abstract

Batch effects are recognized as major sources of technical confounding in high-throughput assays. However, their impact on organoid studies receives little attention in the literature. As organoids gain prominence as a class of emerging new approach methodologies (NAMs), consideration of batch variation will become increasingly important to ensure data reproducibility and accurate interpretation in pre-clinical and clinical studies. In this manuscript, we provide a practical description of our work in detecting, characterizing, and correcting batch effects in a prior published retrospective clinical colorectal cancer organoid drug-response study. We outline the workflow we employed, including exploratory diagnostics, experimental drift detection, and statistical adjustment. We detail the methods employed to evaluate batch effects, monitor longitudinal drift, and select approaches to remove technical artifacts, preserve biological signal and test for robustness. Our experience demonstrates that in even modestly sized studies, results can be adversely affected by insufficient consideration and attempts at ameliorating batch effects. By documenting the challenges we encountered and the solutions implemented within our study, we hope that we can provide a seminal practical reference for organoid researchers and enable increased discussion and adoption of robust batch-compensation practices in the organoid field, ensuring that the topic is more routinely addressed, improved, and eventually standardized.

## Introduction

Organoids are self-organizing 3D multicellular culture systems arising from stem cells ^1^, or in the case of patient derived tumor organoids (PDTOs), from dissociated tumor cells ^2^. Organoids possess the ability to mimic the genetics, molecular characteristics and physiological behaviors of their tissue of origin and thus promise biomimicry exceeding traditional cell-line or animal models ^2^. Organoids and their various applications have seen a surge of interest in recent years. Since Sato *et al.* successfully developed crypt-villus organoids from Lgr5+ intestinal stem cells in 2009 ^3^, the modern organoid field has undergone explosive growth. Following the crypt-villus organoids, many other tissue types have been successfully developed including pancreas, lung, liver, skin, heart, stomach and colon ^4^. Not only have the available tissues expanded, but organoids have experienced ever-broadening applicability. Organoids are routinely used in the fields of regenerative medicine, reproductive biology, infectious, neurodegenerative and genetic disease research, and oncology ^5^. Within the field of oncology, their clinical relevance is becoming increasingly prominent ^6^. Studies have demonstrated the potential of organoids to significantly correlate with or predict patient treatment response in malignancies including colorectal cancer, gastroesophageal cancer and ovarian cancer, and have reported encouraging correlations in a wide selection of other tumor types ^6^. Furthermore, since 2014, there have been a growing number of clinical trials in oncology, utilizing organoids for purposes including the assessment of genetic makeup and drug sensitivity compared to the original tissue, testing of new treatments and establishment of biobanks ^2^. Existing large scale organoid biobanks including the Hubrecht Institute and Hub, promise to make organoids more broadly accessible and standardized for commercial and academic initiatives ^7^. In the wake of the FDA Modernization Act 2.0 in 2022 ^8^ and the the movement away from animal-only studies by the NIH ^9^, as well as the foundation of the NIH Standardized Organoid Modeling (SOM) Center ^9^ in 2025, the organoid field continues to see widening interest and the investigation and utilization of organoids and related technologies are certain to continue to grow in prominence. While the existing impact and untapped potential of organoid technology are clear, challenges unquestionably remain before that their utility can be fully realized ^1^. Tackling the problems of standardization, reproducibility, and reliability will all be core to the successful widespread adoption and realized benefit of organoid technology ^10^. Differences in cell procurement, sample storage, media composition and downstream experimental protocols are only a few examples of considerations that have the potential to introduce variability sufficient to render ostensibly congruous experiments incomparable ^7^. The introduction of standards, shared protocols and industrial developments including automation and miniaturization will be at the core of addressing such challenges ^2,4,6,11,12^. Mature analysis protocols will also be needed to ensure robust data analysis that circumvents analytical pitfalls and produces accurate, biologically meaningful and interpretable outputs from organoid-based experiments and single or multi-site studies.

The phenomenon of batch effects is well known and described in the scientific literature ^13^. These effects are broadly defined as variability introduced to scientific measurements by technical factors that are irrelevant to the hypothesized driver of the experimentally assessed signal. This systematic variability can vary in cause and prominence and depending on its underlying cause it may serve to either attenuate the actual signal being measured or alternatively create the illusion of signal correlated with experimental variables of interest where none actually exists ^14^. When this occurs, it is said that the signal of interest has been confounded by the unwanted technical factor, or confounder ^15^. When the biological variable of interest becomes impossible to differentiate from technical factors due to inadequate experimental design causing them to become inseparably correlated, the experiment can be said to be affected by perfect confounding ^16^. While consideration of batch effects in the literature appears heavily skewed toward high-dimensionality data, awareness of batch effects is necessary in any area of science when samples of any nature are being gathered and measurements of any class are being collected and compared. An historical example that illustrates this point was provided by WJ Youden in 1972 and describes how estimates of Astronomical Unit collected by physicists at different times, in different laboratories, varied both within and across labs and differed from the accepted value^17^.

While there has been documented awareness of nuisance factors or batch effects dating from the early 20th century ^18^, the advent of microarrays and broad-scale multicenter studies in the late 1990s and early 2000s brought about a renaissance in their consideration ^19^. During this period multiple high profile scientific studies saw their conclusions thrown into disarray by the batch-aware third-party analysis ^20^. This led to retraction or damaged credibility for studies reporting genetic differences in ethnic groupings ^21,22^ and genetic variants associated with exceptional longevity in humans ^23,24^ among other examples. The highlighting of these issues on a broad scientific stage had the benefit not only of preventing erroneous scientific conclusions being cemented in common belief, but also of spurring the creation of methodologies, initiatives and standards for their consideration and control ^25–27^.

In an era of multiomics and single cell-level analyses, the awareness of batch effects in the scientific literature appears higher than ever before. Since 2020 there have been over 150 published manuscripts incorporating the term batch effects in their title, which is more than in the 30 years preceding. Despite this previously unseen prominence of literature considering the dangers of batch effects in modern science, and the parallel increase in organoid literature, the two topics rarely intersect. To our knowledge there is no published literature that considers batch effects in organoid studies either critically or in depth. There is undoubtedly awareness of the variability inherent to organoid studies. Published works discuss batch-to-batch variability with specific mention of challenges including extracellular culture matrices ^4,10,28–31^, manual handling ^11^, media variability ^4,31^, clonal differentiation ^31,32^, organoid morphology and gene expression ^12^, and stability of drug sensitivity ^33,34^. While some advice is given toward attempting avoidance of variability upfront, published works discussing analytical methods of control or correction are seemingly absent from modern scientific literature. When analytical methods are discussed they tend to focus upon on-plate microenvironmental conditions most associated with high-throughput screens, while inter-plate variability tends to mainly consider vehicle and kill controls for dose response curve normalization ^35^, rather than experimental confounding. Manuscripts detailing PDTO-based clinical correlation studies appear to make little or no mention of batch effects or their control. An example of the paucity of attention given to batch-effects is a recent systematic review of clinical correlation in colorectal cancer organoid studies ^36^. The review itself made no mention of batch effects, nor did the twenty individual studies described within it mention batch effects as considerations in their experimental design or analysis. The lack of described handling of the issue is perhaps partially due to the fact that the majority of recent published literature on the topic of batch effect handling and correction has been heavily biased toward high-throughput data modalities ^13,14,37,38^. However, preparedness for batch effects in organoid studies is vitally important, particularly as the organoids undergo such explosive growth in use and edge ever closer to status as a clinical technology.

We recently described a collaborative study with the German Cancer Research Center (DKFZ) ^39^, utilizing our micro-fluidics based MOSGen technology ^40,41^ as the foundation for a retrospective clinical correlation study in colorectal cancer patients treated with neoadjuvant standard-of-care agents. Patient-derived tumor tissue was distributed within 3D hydrogel spheres termed Microorganospheres (MOS) with size, shape and consistency under a high degree of control from mature, automated equipment and protocols. The study outcomes demonstrated clinical correlation between patient clinical outcomes and drug-treated MOS models. Despite benefiting from mature automation and state of the art micro-fluidics technology, our study was nonetheless designed from the outset to monitor for batch effects and was able to detect and correct for effects that would otherwise have caused negative impacts on clinical correlation analysis. In this manuscript we detail the analytical steps employed for detection, the nature of the effects observed and the approaches taken to correct them and ensure robustness of the post-correction results. The relatively small scale of an organoid dataset compared to high-dimensionality omics and multiomics data, and the disparate structure of the data itself meant that deviations from methodologies considered standard in other fields were required. It is our belief that effects similar to or more severe than those we observed are of an even higher likelihood in studies that lack the standardized automated components that we have developed, and we hope that the work we describe will inform others in the field to maximize the integrity of results from their future organoid studies.

### Study Recap

The study is described fully in our recently published manuscript ^39^ but in brief summary: Primary or metastatic lesions were collected from colorectal cancer patients that had been treated clinically with neoadjuvant standard-of-care therapies. Organoids were established from patient tumor samples and deposited in MOS droplets following sample dissociation. MOS droplets were deposited at an average density of 40 per well in 384 well plates, drug-treated with a nine-point dose gradient and dose response (negative logarithm of the half maximal inhibitory concentration i.e. pIC50) calculated using fluorescence intensity from brightfield microscopy with EpCAM staining, following 7 days in-assay. EpCAM signal intensity at assay day 0 was used to normalize EpCAM endpoint signals to account for minor differences in starting biomass. Percentiles of response to each standard-of-care therapy were calculated and utilized in subsequent clinical correlation analyses. Clinical correlation was based on percentile of in-assay MOS model dose response to the patient’s standard of care treatment, based on both binary clinical lesion-level clinical response (radiological RECIST-like scoring for metastatic tumors and pathological Dworak score for primary lesions), and disease-free survival (DFS). Endpoint CellTiter-Glo assays were run in parallel to EpCAM imaging as an orthogonal measurement of MOS model viability.

### General experimental design

Besides analytics to monitor or post-correct batch effects, core good practices are vital in experimental design. Post-hoc correction of batch effects is not always possible, nor is it necessarily straightforward. There is no substitute for careful planning and good experimental design. We will not describe all elements of good design in depth since they are covered comprehensively elsewhere ^15,19^, but we will give brief consideration to a selection of some vital design considerations and how they were approached in our study.

1) Randomization Ensuring that samples representing the experimental variables of interest are well randomized across time, instruments, etc. is important to avoid high levels of confounding. For example, clustering most responder samples in one month and most non-responders in the following month introduces the potential of observing differences in experimental measurements that can be mistaken as being due to response status, while they are in fact due to temporal differences that may be linked to various underlying causative phenomena e.g. a change in machine calibration, or media batch. Clinical studies like our own can be inherently problematic, since sample receipt and processing is frequently dictated by a patient’s disease progression and clinical scheduling. Despite being unable to formally randomize conditions, we were able to ensure a relatively randomized order of sample processing as shown in Table 1.

**Table 1:** Clinical and experimental metadata for all samples (n=37) included in clinical correlation analysis.

| Sample | Study Phase | Processing Batch | Tumor-level clinical response | Clinical treatment | Media Batch | Localization | Disease free survival (years) |
| --- | --- | --- | --- | --- | --- | --- | --- |
| Sample 1 | Phase 1 | 1.1 | Progression | FOLFOX | NA | Lymph node | 0.76 |
| Sample 2 | Phase 1 | 1.1 | Response | FOLFIRI | NA | Liver metastasis | 0.58 |
| Sample 3 | Phase 1 | 1.1 | Response | FOLFOX | NA | Primary tumor | 1.95 |
| Sample 4 | Phase 1 | 1.2 | Response | FOLFOX | NA | Liver metastasis | 0.2 |
| Sample 5 | Phase 1 | 1.2 | Response | FOLFOX | NA | Primary tumor | 0.54 |
| Sample 6 | Phase 1 | 1.2 | Progression | FOLFOX | NA | Primary tumor | 3.41 |
| Sample 7 | Phase 1 | 1.3 | Response | FOLFOX | NA | Liver metastasis | 0.3 |
| Sample 8 | Phase 1 | 1.3 | Response | FOLFOX | NA | Liver metastasis | 1.06 |
| Sample 9 | Phase 1 | 1.4 | Response | FOLFOX | NA | Liver metastasis | 0.3 |
| Sample 10 | Phase 1 | 1.4 | Response | FOLFOX | NA | Liver metastasis | 1.06 |
| Sample 11 | Phase 1 | 1.4 | Response | FOLFOX | NA | Liver metastasis | 1.11 |
| Sample 12 | Phase 2 | 2.1 | Response | FOLFOX | 1 | Primary tumor | 3.42 |
| Sample 13 | Phase 2 | 2.1 | Response | FOLFOX | 1 | Primary tumor | 0.33 |
| Sample 14 | Phase 2 | 2.1 | Progression | FOLFOXIRI | 1 | Primary tumor | 0.71 |
| Sample 15 | Phase 2 | 2.1 | Response | FOLFOXIRI | 1 | Primary tumor | 0.81 |
| Sample 16 | Phase 2 | 2.1 | Response | FOLFOX | 1 | Liver metastasis | 1.68 |
| Sample 17 | Phase 2 | 2.1 | Response | FOLFOX | 1 | Primary tumor | 0.25 |
| Sample 18 | Phase 2 | 2.1 | Response | FOLFOX | 1 | Liver metastasis | 0.54 |
| Sample 19 | Phase 2 | 2.2 | Progression | FOLFOX | 2 | Liver metastasis | 0.33 |
| Sample 20 | Phase 2 | 2.2 | Response | FOLFOXIRI | 2 | Liver metastasis | 0.71 |
| Sample 21 | Phase 2 | 2.2 | Response | FOLFOXIRI | 2 | Liver metastasis | 0.71 |
| Sample 22 | Phase 2 | 2.2 | Response | FOLFOXIRI | 2 | Liver metastasis | 0.81 |
| Sample 23 | Phase 2 | 2.2 | Response | FOLFOX | 2 | Liver metastasis | 1.11 |
| Sample 24 | Phase 2 | 2.3 | Response | FOLFIRI | 3 | Liver metastasis | 0.04 |
| Sample 25 | Phase 2 | 2.3 | Response | FOLFOXIRI | 3 | Liver metastasis | 0.81 |
| Sample 26 | Phase 2 | 2.4 | Response | FOLFOX | 3 | Liver metastasis | 2.63 |
| Sample 27 | Phase 2 | 2.4 | Response | FOLFOX | 3 | Liver metastasis | 2.63 |
| Sample 28 | Phase 3 | 3.1 | Progression | FOLFOX | 4 | Liver metastasis | 0.33 |
| Sample 29 | Phase 3 | 3.1 | Response | FOLFIRI | 4 | Liver metastasis | 0.45 |
| Sample 30 | Phase 3 | 3.2 | Progression | FOLFOX | 5 | Liver metastasis | 0.33 |
| Sample 31 | Phase 3 | 3.2 | Progression | FOLFOX | 5 | Liver metastasis | 0.33 |
| Sample 32 | Phase 3 | 3.3 | Response | FOLFOX | 5 | Lymph node | 0.35 |
| Sample 33 | Phase 3 | 3.3 | Progression | FOLFOX | 5 | Liver metastasis | 0.33 |
| Sample 34 | Phase 3 | 3.4 | Progression | FOLFIRI | 6 | Primary tumor | 0.02 |
| Sample 35 | Phase 3 | 3.4 | Progression | FOLFIRI | 6 | Liver metastasis | 0.02 |
| Sample 36 | Phase 3 | 3.5 | Response | FOLFOX | 6 | Liver metastasis | 0.35 |
| Sample 37 | Phase 3 | 3.5 | Progression | FOLFIRI | 6 | Primary tumor | 0.02 |

2) Avoid perfect confounding This is related to the previous point but deserves specific mention - perfect confounding should be avoided at all costs. In a study like ours where drug response is the variable of interest, it is key to ensure that clinical responders and non-responders are sufficiently randomized across potential confounding conditions to ensure that should batch effects occur, the opportunity exists to attempt disambiguation of a variable of interest and a confounder. In the case of our study, while non-responders were less frequently observed than responders, they were effectively distributed across the timeline of the study to avoid over-clustering that might have been problematic. Nonetheless it was not possible to wholly avoid batches containing only one response group of interest, nor was it possible to perfectly balance the number of responders and non-responders within or between batches.
3) Standardized operating procedures Ensuring that sample procurement, handling and processing procedures are well documented and standardized, with qualified operators trained in their use is important in reducing technical variability that can affect a study. Within both the clinical setting and in our laboratory, standard procedures were frequently in place and relevant individuals were appropriately trained to follow them. Nonetheless, the study was conducted in a non-operations environment while the laboratory was early in its lifecycle and thus some protocols were in flux, meaning potential for inconsistencies existed.
4) Automate processes where possible Automation has the potential to reduce manual error and variability throughout an experiment and can thus attenuate specific batch effects caused by inaccuracies, or inconsistencies introduced by operators or manually operated equipment. Our experimental setup was widely automated. The MOSgen apparatus automated and controlled cell deposition and MOS droplet formation. Furthermore, automated liquid handling was employed for MOS droplet deposition in experimental plates, addition of fluorophores and drug dosing.
5) Avoid changing procedures during a study As much as possible, procedures, methodologies, reagents etc should be kept as constant as possible through a study. In the context of our study, formal change control procedures were followed. Material changes to experiments were made as infrequently as possible, were discussed by a multidisciplinary team prior to implementation, and impact assessments made before changes believed to be acceptable were implemented and documented.
6) Maintain records of possible confounders It is not always possible to be aware of every confounder that might affect an experiment, but prior to a study taking place, potential confounders should be discussed and documented. Furthermore, a record of these should be kept for every sample processed to enable post-hoc analysis of potential confounding features, and correction if possible. In the case of our study, potential confounders including media batch, processing batch, etc were widely recorded and ultimately empowered downstream analysis when the emergence of batch effects was suspected.
7) Include bridging samples or similar technical replicate controls A recommended approach to monitoring and potentially correcting for batch effects is inclusion of a control sample which is run in every experimental batch and whose results are expected to be as close as possible to identical with every run. This can expedite the detection of batch effects since any marked deviation in its results are an obvious sign of an issue, and in some cases the sample itself can be used to calculate a corrective factor that can be used to normalize the other samples in the batch. While our experiments did utilize bridge samples, due to a combination of logistical factors, they were not always available for every batch and drug of interest. Thus, while they could provide us with some indication of batch effects, they were insufficient in isolation to correct those effects.

### On-plate control measures and study monitoring

Platemaps were used to standardize plate layout and drug dosing. Edge wells were used as buffer rows and platemaps were designed in order to avoid spatial biases. All doses and conditions were run in quadruplicate. In all cases, dose response curves were normalized to multiple control wells (zero killing) and staurosporine kill wells (full killing). After laboratory processing of each sample batch, dose response metrics were calculated and plotted in the absence of clinical response information to enable monitoring of undesired trends or drift. The study was performed in 3 temporal phases (referred to as Phase 1-3). Exploratory analysis incorporating clinical response data was conducted at the conclusion of each phase to assist in detection of potential batch effects.

### Cohort Details and Metadata

Full cohort details are provided in Table 1. A total of 37 neoadjuvant treated colorectal cancer samples from 21 individual patients passed QC and were included in the final clinical correlation analysis. The three study phases comprised a total of thirteen processing batches. Patients were treated clinically with FOLFOX, FOLFIRI or FOLFOXIRI. Samples included primary tumor, liver metastases or lymph node metastases. Primary location was classified as either rectal, or left or right colon. Disease free survival time ranged from 0.02 to 3.42 years. Eleven samples were clinically classified as non-responsive to treatment, while 26 were classified as responsive. Processing batch metadata was available for all samples. Media batch was recorded for study Phases 2-3 only. Drug batch was not recorded since a policy of drug discardment after three freeze-thaw cycles was adopted and this practice was believed to be sufficiently rigorous to avoid drug-related issues within the experiment.

### Longitudinal trend analysis

Log logistic dose response curves were generated for all sample-treatment combinations and pIC50 values were calculated as a measure of overall sample sensitivity to a treatment. pIC50s for each individual sample-treatment combination were plotted temporally, post-processing of each batch, to serve as an indicator of potential trends (Figure 1A). A 3-sample rolling mean was plotted to aid with the identification of temporal trends amidst expected natural variability in pIC50 values. Rolling-mean smoothing is widely used in identifying temporal trends and detecting drift across many scientific and applied fields that include finance ^42^, signal processing ^43^, climate science ^44^, epidemiology ^45^, business forecasting ^46^, and standard time-series analysis ^47^. Although rolling-window methods can be sensitive to irregular time spacing, in this study the elapsed time between runs had no expected biological or technical effect. Therefore, sequence order rather than absolute time was the relevant axis for drift detection.

**Figure 1A:**
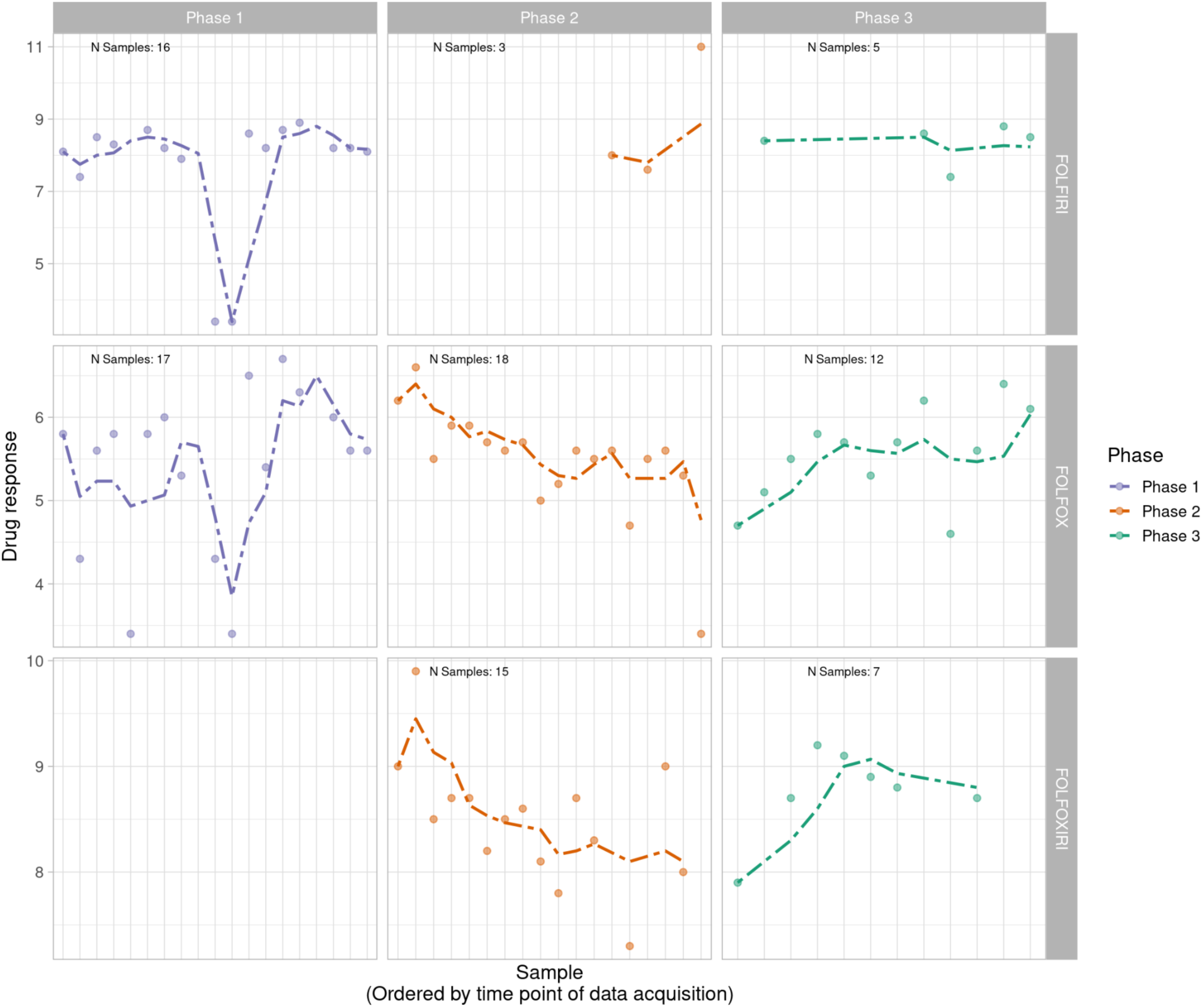
Longitudinal monitoring of patient-derived MOS model drug response values (pIC50) across all phases of the project timeline with phases (Phase 1, 2 and 3) displayed from left to right and treatment type (FOLFIRI, FOLFOX, FOLFOXIRI) displayed from top to bottom. Dotted lines display a right-aligned 3-sample window rolling mean pIC50 value to account for fluctuations and aid with trend identification. Phase 1 does not appear to show a clear trend while FOLFOX and FOLFIRI show potential downward and upward trends in Phase 2 and Phase 3 respectively.

While Phase 1 data revealed minimal visual trends, Phases 2 and 3 appeared to indicate potential linear trends with FOLFOX and FOLFOXIRI data suggestive of a downward trend in Phase 2 (decreasingly responsive) and an upward trend in Phase 3 (increasingly responsive). FOLFIRI in Phase 2 suggested an upward trend but was heavily influenced by a small number of samples and a single outlier, while FOLFIRI in Phase 3 appeared flat. FOLFOXIRI was not included in Phase 1 of the study.

While this analysis in isolation was insufficient to conclude a problem, it was nonetheless a valuable representation of the project timeline that indicated potential patterns warranting further investigation. Since endpoint CellTiter-Glo assays were run in parallel to EpCAM imaging, we were able to determine that similar longitudinal patterns existed using this orthogonal measurement of viability. This enabled us to conclude that any trend being observed was independent of EpCAM batch, brightfield microscopy conditions, or related variables.

Principal components analysis was attempted due to its traditional use in batch effect exploration, but was unrevealing, likely due to the limited scale and dimensionality of the dataset, and limited batch sizes (data not shown).

### Linear modeling

Linear modeling was performed on a per-drug, per-phase basis to follow up robustly on visual trend inspection and determine if any recorded variables showed significant correlation with the MOS model drug responses. Variables that were considered included processing batch, media batch, clinical resistance status, and DFS. Phase 1 showed statistically significant correlation with clinical tumor responses for both FOLFOX and FOLFIRI (p=0.007 and p=0.0007 respectively). Phase 2 demonstrated statistically significant or close to significant correlation with an assortment of processing batches and media batches for FOLFOX, FOLFIRI, and FOLFOXIRI (p-values ranging from 0.011 to 0.103). Phase 3 also primarily showed correlations with media and processing batches for all drugs, although not statistically significant. On the basis of this analysis we concluded that, as suggested by trend visualization, Phase 1 was likely effectively free from batch effects while Phases 2 and 3 were each impacted to varying degrees. Both media batch and processing batch appeared relevant to latter stage batch effects, and since these frequently varied alongside one another and media batch was incompletely captured, processing batch was identified as the likely optimal surrogate metadata field to inform attempted batch compensation.

### Batch effect analysis

Batch compensation was subsequently investigated using the ComBat function from the sva package in R ^48^. While ComBat’s origins lie in microarray studies, it has shown versatility in broader settings and smaller datasets ^49–52^, and has been recommended for drug response studies ^53^. Batch variables suspected to affect experimental measurements are passed to the function using the provided ‘batch’ argument, with processing batch used in our case. Output is a set of corrected measurements with batch effects removed, where downstream analysis techniques can subsequently be applied to the data. Removal of batch effects and use of surrogate variables has been demonstrated to improve reproducibility, stabilize error rate estimates and reduce dependence ^48^.

We implemented the ComBat in mean-only mode which applies mean centering across batches without variance/scale adjustment. In this mode, batch effects are modeled as additive offsets and removed through estimation of batch-specific parameters. The mean-only option was chosen because the dataset had low dimensionality, and heterogeneous variance was expected across batches due to variable and unpredictable drug responses ^54^. This approach reduces systematic batch effects while preserving within-batch variance that could reflect true biological differences. Responder status was supplied to ComBat as a biological covariate to prevent overcorrection and maintain response-group-related relevant signals, as has been recommended by the original authors and elsewhere ^48,55^. ComBat builds a linear model on batch and biological covariates to estimate additive batch offsets. When mean-only mode is selected, the implementation subtracts only those offsets, leaving scale and variance untouched ^56^.

Post-ComBat batch compensation, dose response data was replotted longitudinally and visualized in the absence of clinical correlation data. The trends suggested by the original rolling mean visualization appeared attenuated by ComBat, while Phase 1 results appeared relatively unaffected (Figure 1B).

**Figure 1B:**
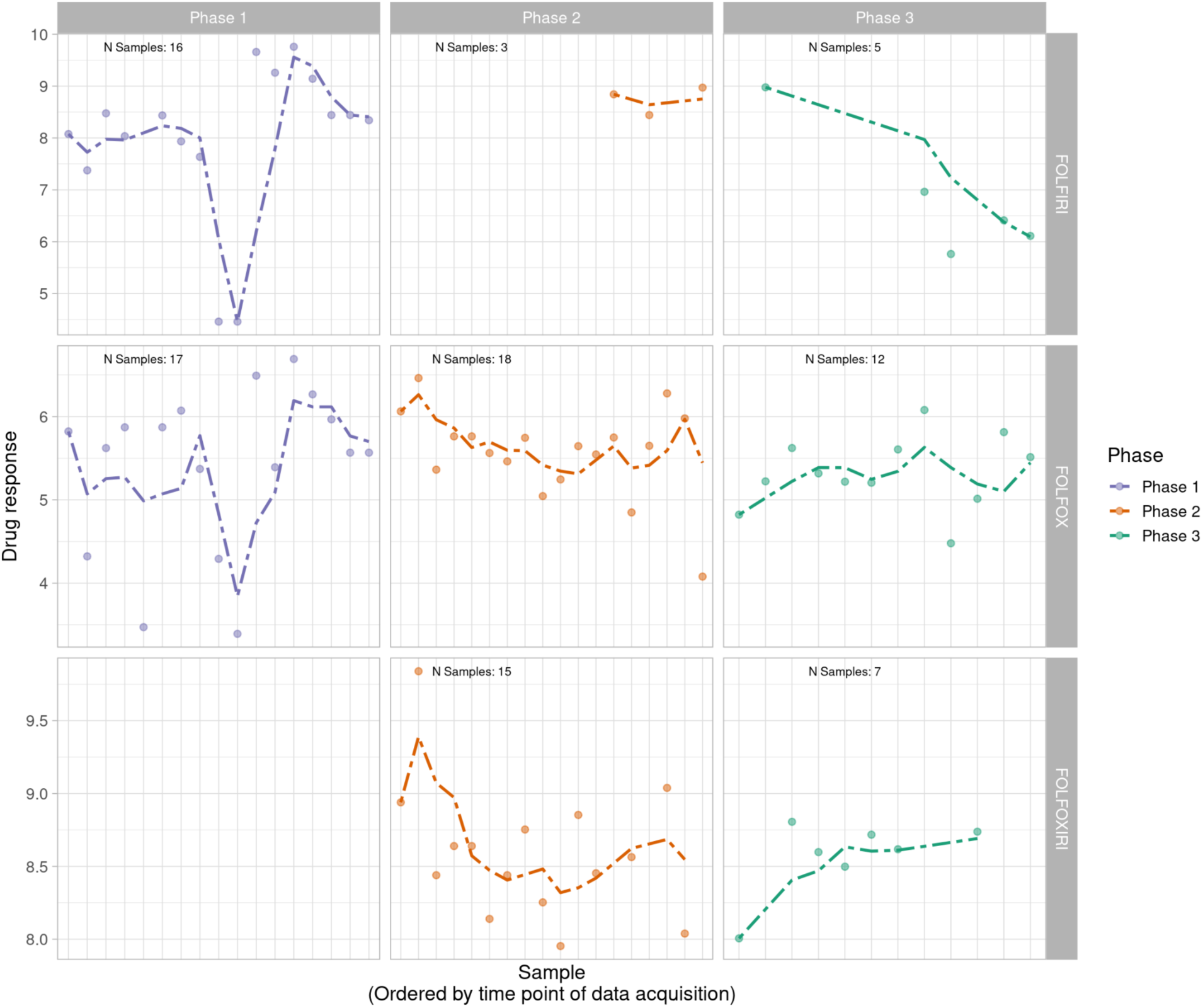
Longitudinal monitoring of post-ComBat patient-derived MOS model drug response values (pIC50) across all phases of the project timeline with phases (Phase 1, 2 and 3) displayed from left to right and treatment type (FOLFIRI, FOLFOX, FOLFOXIRI) displayed from top to bottom. Dotted lines display a right-aligned 3-sample window rolling mean pIC50 value to account for fluctuations and aid with trend identification. Trends observed with a rolling average prior to ComBat appear visually attenuated after processing by ComBat.

The magnitude and direction of change in pIC50 for each sample post-ComBat was also calculated and visualized per sample (Figure 1C) and per batch (Figure 1D). These results again showed minimal effects on Phase 1 samples, and more pronounced effects in Phases 2 and 3 individually, with some batches showing more modest effects than others. The directions of the observed pIC50 changes per sample were in broad agreement with an expected reversal relative to observed trends in the longitudinal analysis.

**Figure 1C:**
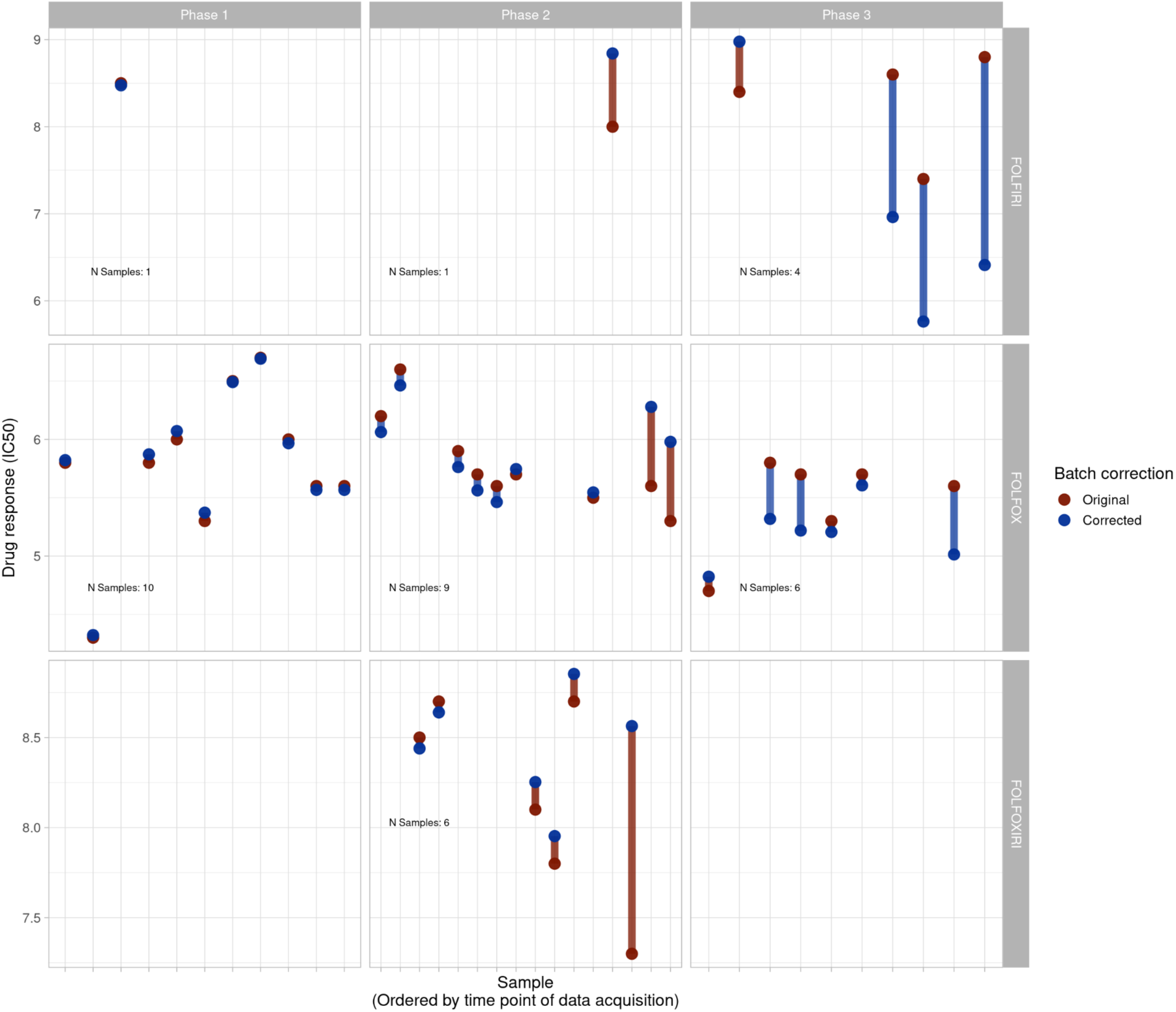
Per-sample change in pIC50 for all samples entering clinical correlation analysis and their clinical treatment only (n=37) pre- and post-ComBat batch compensation across all project phases with phases (Phase 1, 2 and 3) displayed from left to right and treatment type (FOLFIRI, FOLFOX, FOLFOXIRI) displayed from top to bottom. Phase 1 shows minimal shifts while larger changes are visible in Phases 2 and 3, and directionally correspond largely to what would be expected if attenuating the trends observed in longitudinal analysis.

**Figure 1D:**
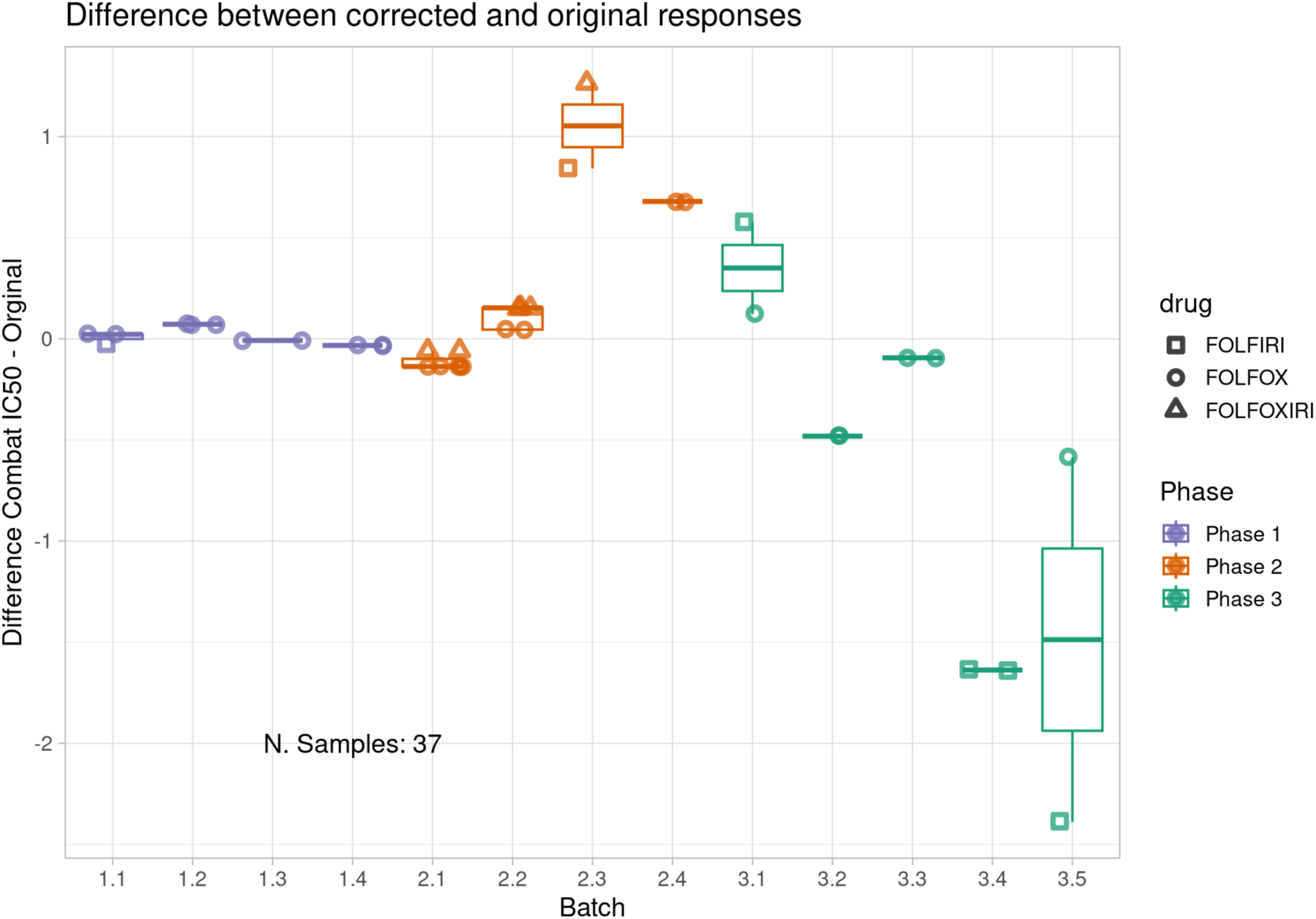
Per batch delta of pre and post-ComBat PIC50 values for all samples entering clinical correlation analysis (and their clinical treatment only (n=37). Batches are displayed on the x-axis while pre and post-ComBat pIC50 delta is displayed on the y-axis. Drug combinations tested in-assay are represented by point shape and color represents study phase. Correction varies by batch but larger deltas are observed in Phases 2 and 3 while Phase 1 shows minimal changes.

### Clinical correlation by project phase

Subsequent to the investigative longitudinal analyses, we plotted boxplots displaying correspondence between patient clinical response and ComBat-compensated patient tumor-derived MOS model dose response. MOS model pIC50 values were expressed as per-drug percentiles to facilitate combination of alternative drugs with disparate potencies in a single analysis. Clear discriminative ability was observed across all study phases, both individually (Figure 2) and combined as a single cohort, as demonstrated in our original study (Figure 3). A cluster-naive Wilcoxon rank sum test was performed for all phases combined and produced a statistically significant p-value of 0.0007. The p-value remained significant following multiple-testing correction using the Benjamini-Hochberg method (p=0.0051). Since the Wilcoxon rank sum test assumes independence of samples and our study included multiple lesions per patient in some cases, we performed a patient-level permutation analysis where response labels underwent permutation for 2000 iterations at the patient level to generate the null distribution. This again yielded a significant p-value (p=0.012). ROC analysis was conducted using the post-ComBat MOS model response values (Figure 4) and yielded an AUC of 0.86 for the full cohort (95%CI: 0.74 - 0.98). For the cutoff on the ROC curve maximizing balanced sensitivity and specificity, the assay showed 83% accuracy (95%CI: 69%-100%), 82% sensitivity (95%CI: 64%- 100%), and 85% specificity (95%CI: 64%-100%).

**Figure 2:**
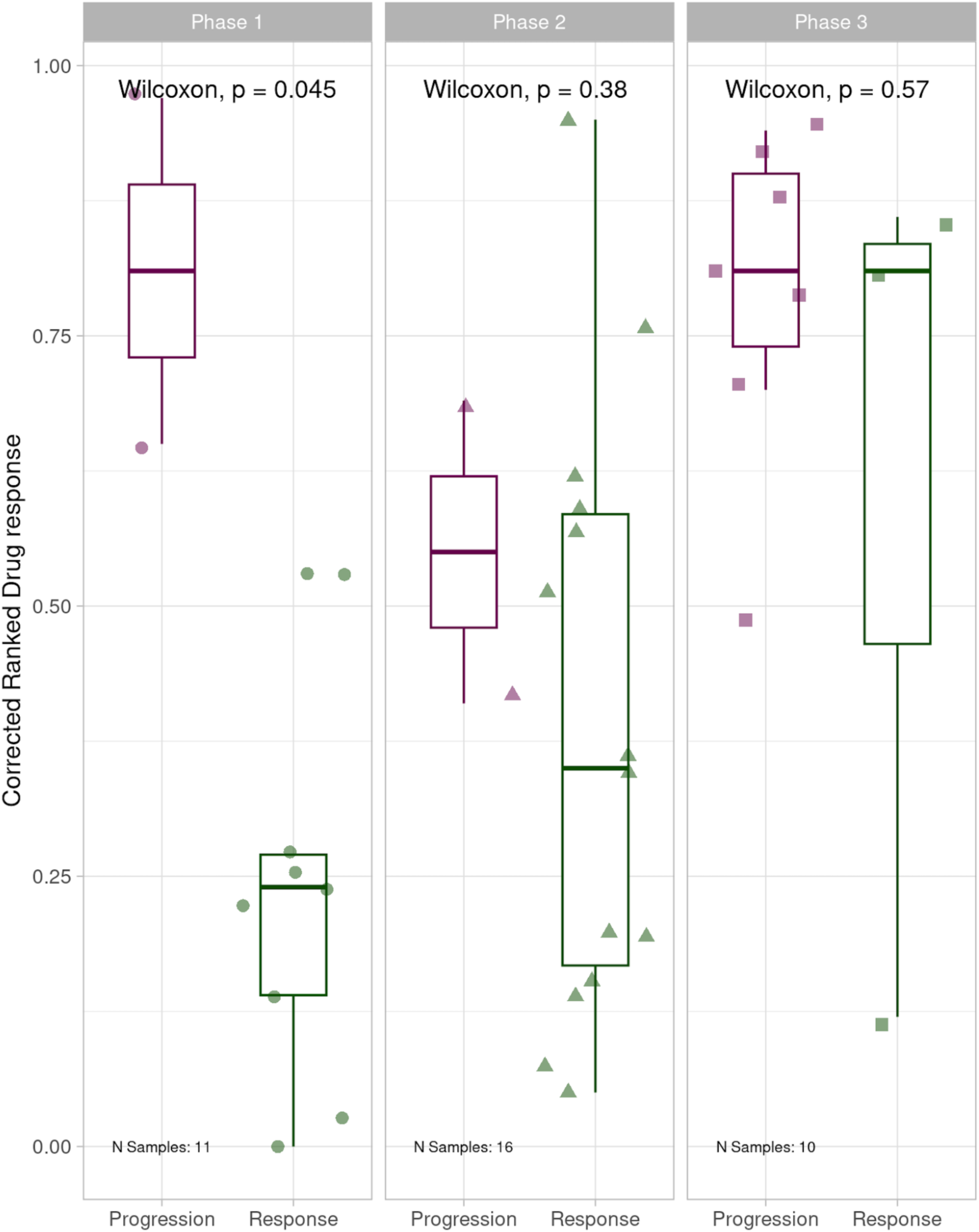
Boxplots displaying correspondence of patient-derived MOS model dose response to clinical treatment response for each project phase, post batch compensation with ComBat. All phases show separation of responders and non responders, with patients responsive to clinical treatment having the most drug-responsive MOS models.

**Figure 3.**
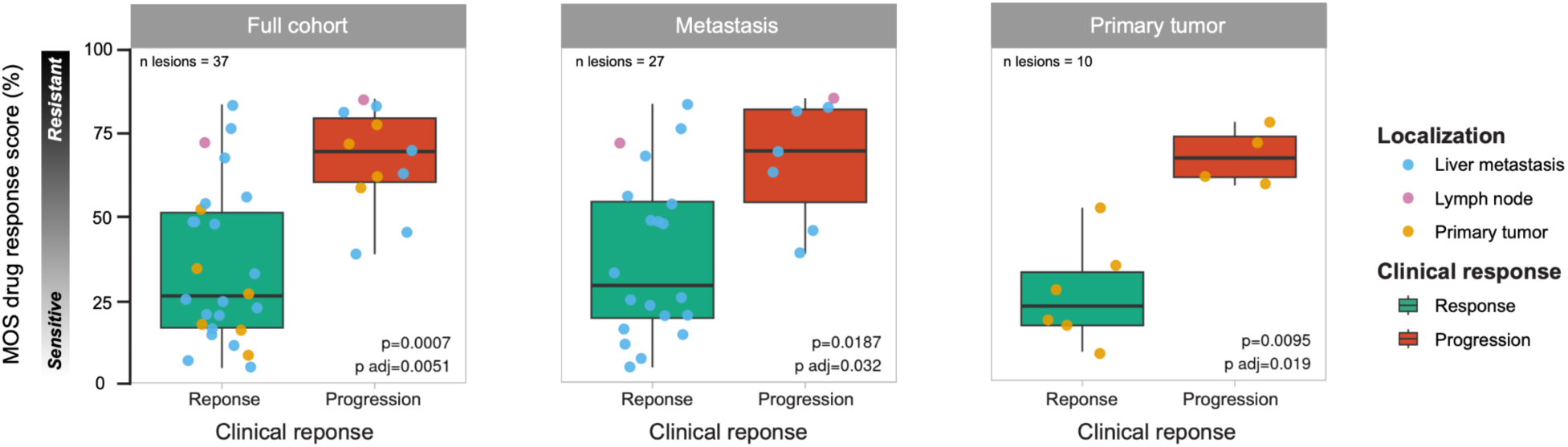
Comparison of MOS drug response score between lesions that showed clinical response or progression in the full cohort (n = 37 lesions from n = 21 patients), metastasis (n = 27 lesions from n = 16 patients), and primary tumor (n = 10 lesions from n = 9 patients) groups. Each point indicates a patient-derived MOS model, and the color indicates tumor origin. Blue = liver metastases, pink = lymph node metastasis, and orange = primary tumor. Statistical significance was assessed using the Wilcoxon rank-sum test, and P values were adjusted for multiple comparisons using the Benjamini–Hochberg method. **Reproduced with permission from Wolters Kluwer Health, Inc. From: Gobits R, Schleußner N, Oliver GR, et al. *Functional Precision Medicine Using MicroOrganoSpheres for Treatment Response Prediction in Advanced Colorectal Cancer*. JCO Precision Oncology. 2026;10(10):e2500501. © American Society of Clinical Oncology.** https://ascopubs.org/doi/10.1200/PO-25-00501 **This figure is reproduced for non-commercial use in this preprint. No data values or analytical content have been modified. This figure is not covered by the Creative Commons license applied to this preprint.**

**Figure 4:**
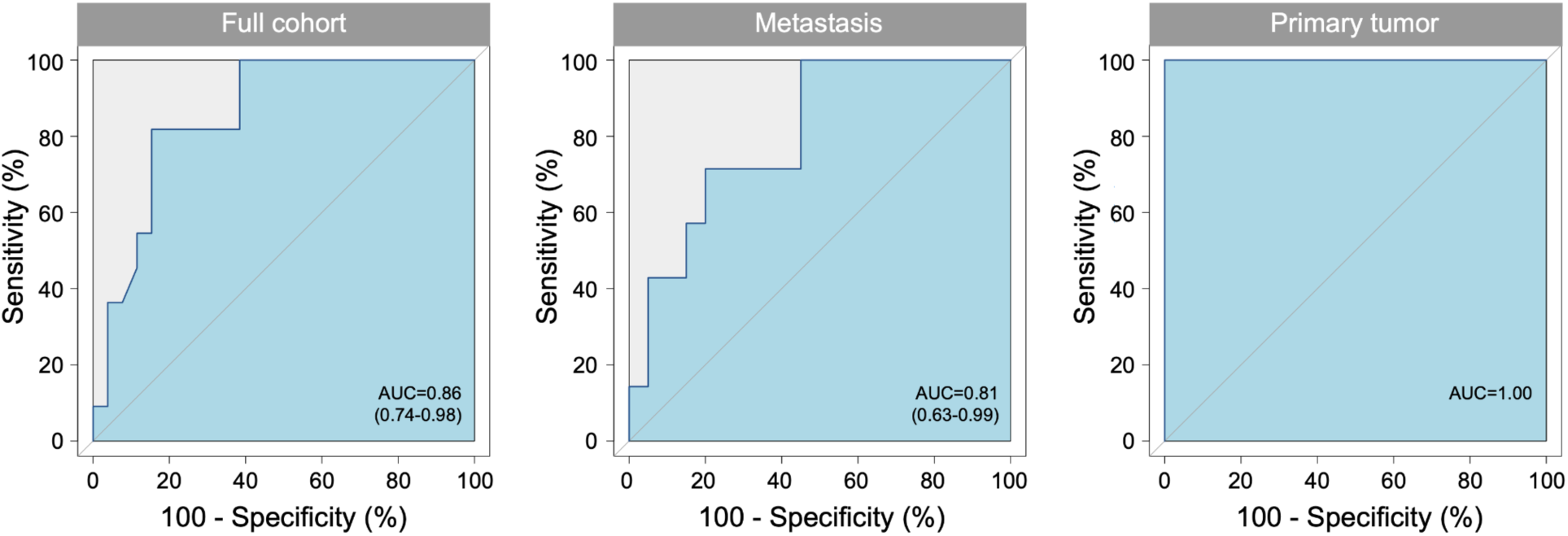
ROC curves for predicting treatment response corresponding to the respective box plots in Figure 3. The curves illustrate the trade-off between sensitivity and specificity across a sliding threshold, with the diagonal line representing an AUCROC of 0.5. AUC is displayed in the plot body, followed by 95% confidence intervals in parentheses. **Reproduced with permission from Wolters Kluwer Health, Inc. From: Gobits R, Schleußner N, Oliver GR, et al. *Functional Precision Medicine Using MicroOrganoSpheres for Treatment Response Prediction in Advanced Colorectal Cancer*. JCO Precision Oncology. 2026;10(10):e2500501. © American Society of Clinical Oncology.** https://ascopubs.org/doi/10.1200/PO-25-00501 **This figure is reproduced for non-commercial use in this preprint. No data values or analytical content have been modified. This figure is not covered by the Creative Commons license applied to this preprint.**

Clinical correlation was subsequently assessed utilizing retrospective binary tumor-level patient clinical treatment response data and corresponding ComBat-*uncompensated* dose response data for drug-treated patient-derived MOS models. Phase 1 demonstrated clean separation of responders and non-responders with more drug-responsive MOS models corresponding to patients’ clinical responsiveness to treatment (cluster naive Wilcoxon rank sum p=0.044). This provided evidence of an initially predictive assay even in the absence of batch-compensation (Figure 5) however the apparent predictivity deteriorated in both Phase 2 (p=0.75) and Phase 3 (p=0.65) of the study with responder and non-responder groupings showing high degrees of overlap and data from all three phases combined similarly showing little distinction between responders and non-responders. Receiver operating characteristic (ROC) analysis for the full study data produced an AUC of 0.421 further highlighting the lack of predictivity (Figure 6). The coincidence of unexpected linear trends in longitudinal monitoring for Phase 2 and 3, accompanied by promising clinical discriminative ability in Phase 1 and subsequent degradation of the assay’s discriminative ability during Phases 2 and 3 provided further indication of likely technical issues impacting the study, as indicated by the original longitudinal analysis.

**Figure 5:**
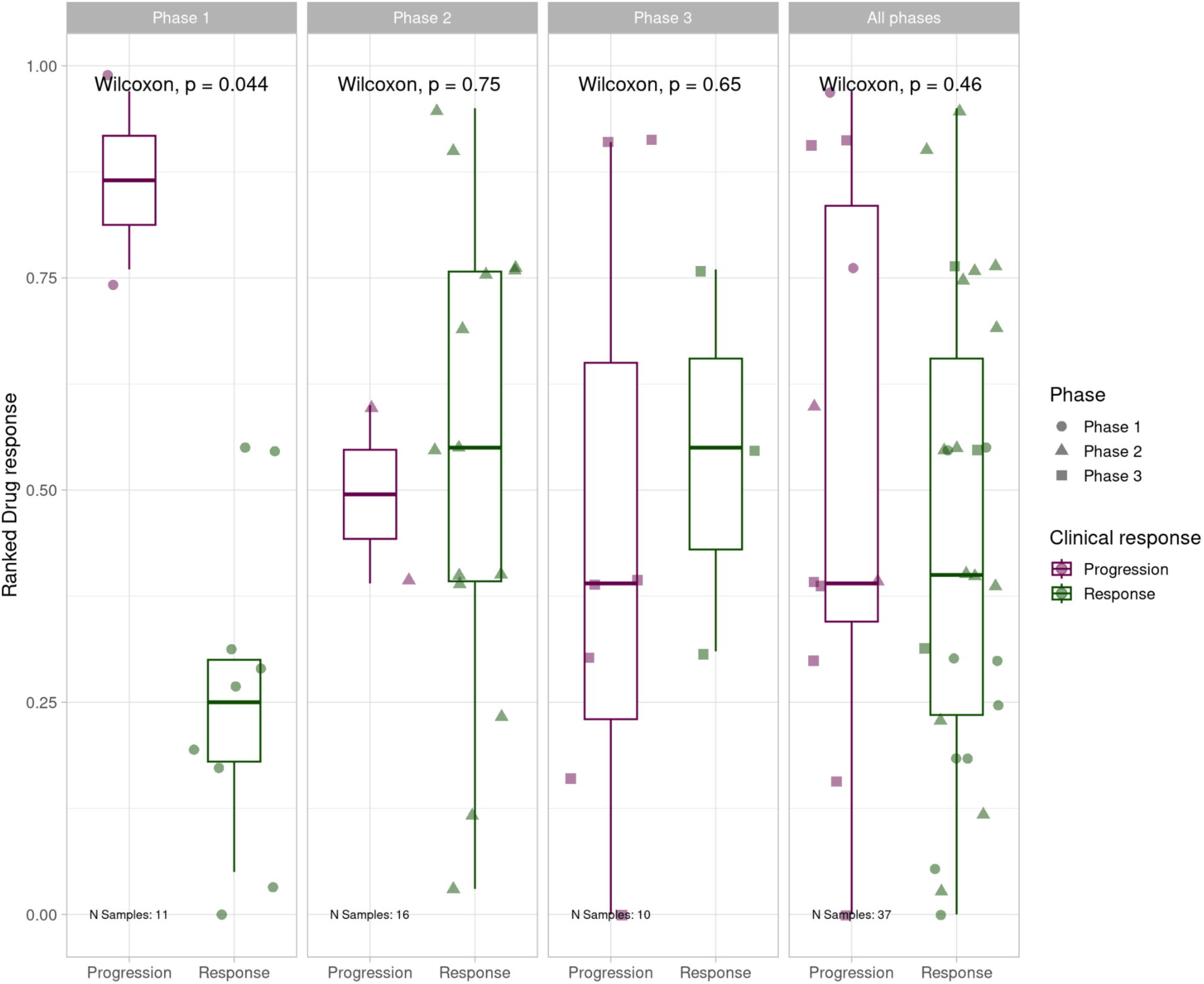
Boxplots displaying correspondence of patient-derived MOS model dose response to clinical treatment response for each project phase individually and in summation, prior to batch consideration. Phase 1 showed clear separation of responders and non responders, with patients responsive to clinical treatment having the most drug-responsive MOS models. Phase 2 and 3 individually, and all phases combined showed no discriminative ability.

**Figure 6:**
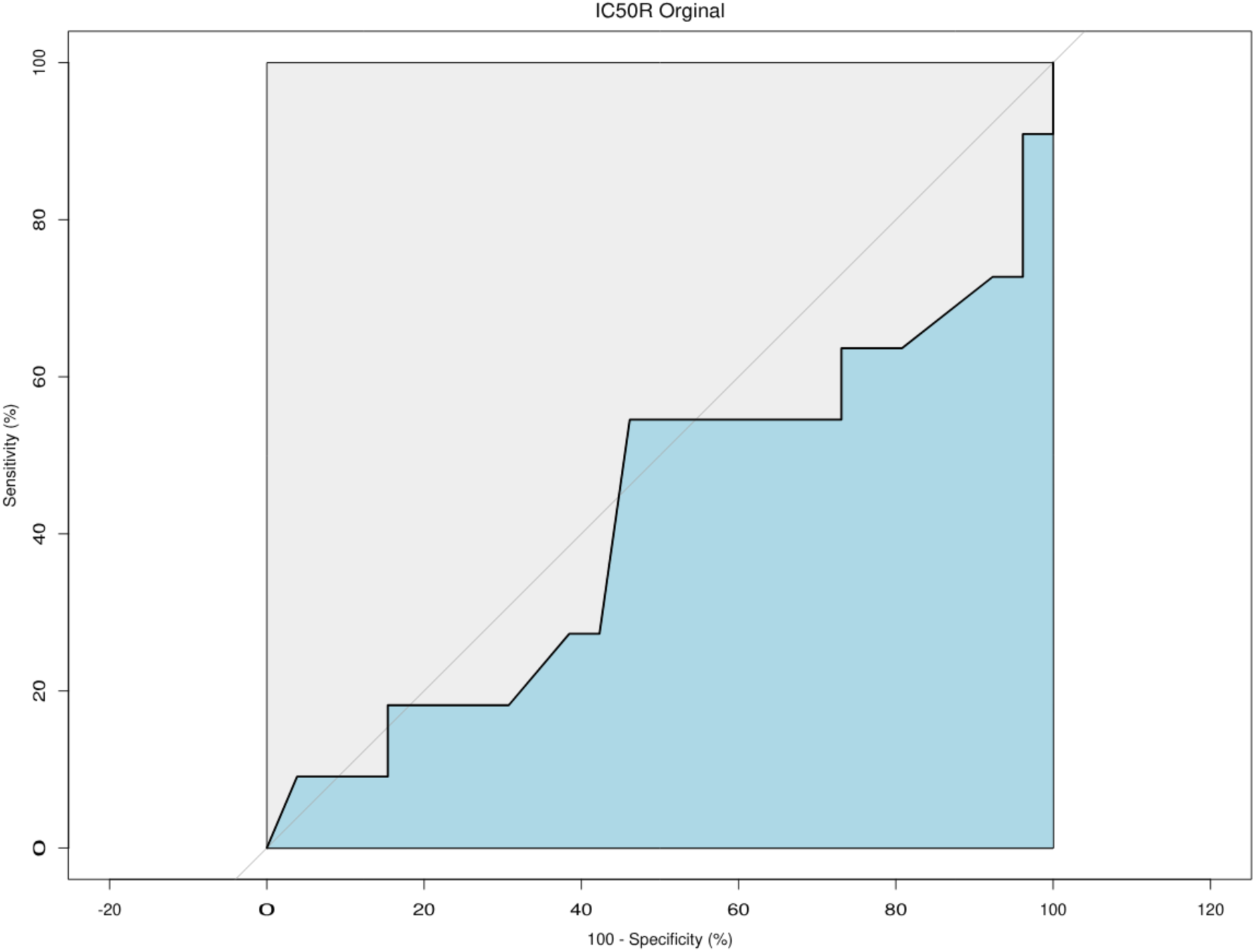
Receiver operating characteristic analysis for all project phases combined showing predictivity of clinical responders (specificity) and non-responders (sensitivity) by patient tumor-derived MOS assay, prior to batch consideration. AUC was 0.421 (95%CI: 0.197-0.645). An AUC of 0.421 reflects predictiveness lower than 0.5 which would be expected by random classification.

### Batch compensation robustness analysis

While post-ComBat outcomes in Phase 2 and 3 matched the clinical discriminative ability observed in pre-ComBat Phase 1 results, and batch compensation appeared to attenuate the longitudinal trends observed across Phases 2 and 3, it remained necessary to confirm the robustness of the post-batch compensation results. Others have questioned the potential for batch effect correction to introduce signals that favor the biological outcome of interest ^57^. These concerns have primarily been identified as affecting high-dimensionality multivariate data with variance-based batch correction and unbalanced batches, and are less likely to apply to a univariate, mean-only correction scenario like ours. Nonetheless, we applied permutation analysis to ensure that genuine compensation of batch differences was driving the correction toward a discriminative outcome and that no artifact of the ComBat analysis could be producing the result. In detail, batch labels were randomly shuffled among samples before the full batch correction process was rerun using the newly labeled samples and the Wilcoxon rank sum test was applied, and effect size calculated. This analysis was repeated for 1000 iterations, generating empirical null distributions. Only one simulation attained a p-value as extreme as our ComBat-compensated analysis (Figure 7A). Furthermore, no iteration of the permutation analysis could produce an outcome with an effect size matching or exceeding that of our original ComBat-compensated analysis (Figure 7B). Collectively these findings indicate that correctly labeled batches were the central driver of the batch correction and no spurious behavior of batch compensation was responsible.

**Figure 7A:**
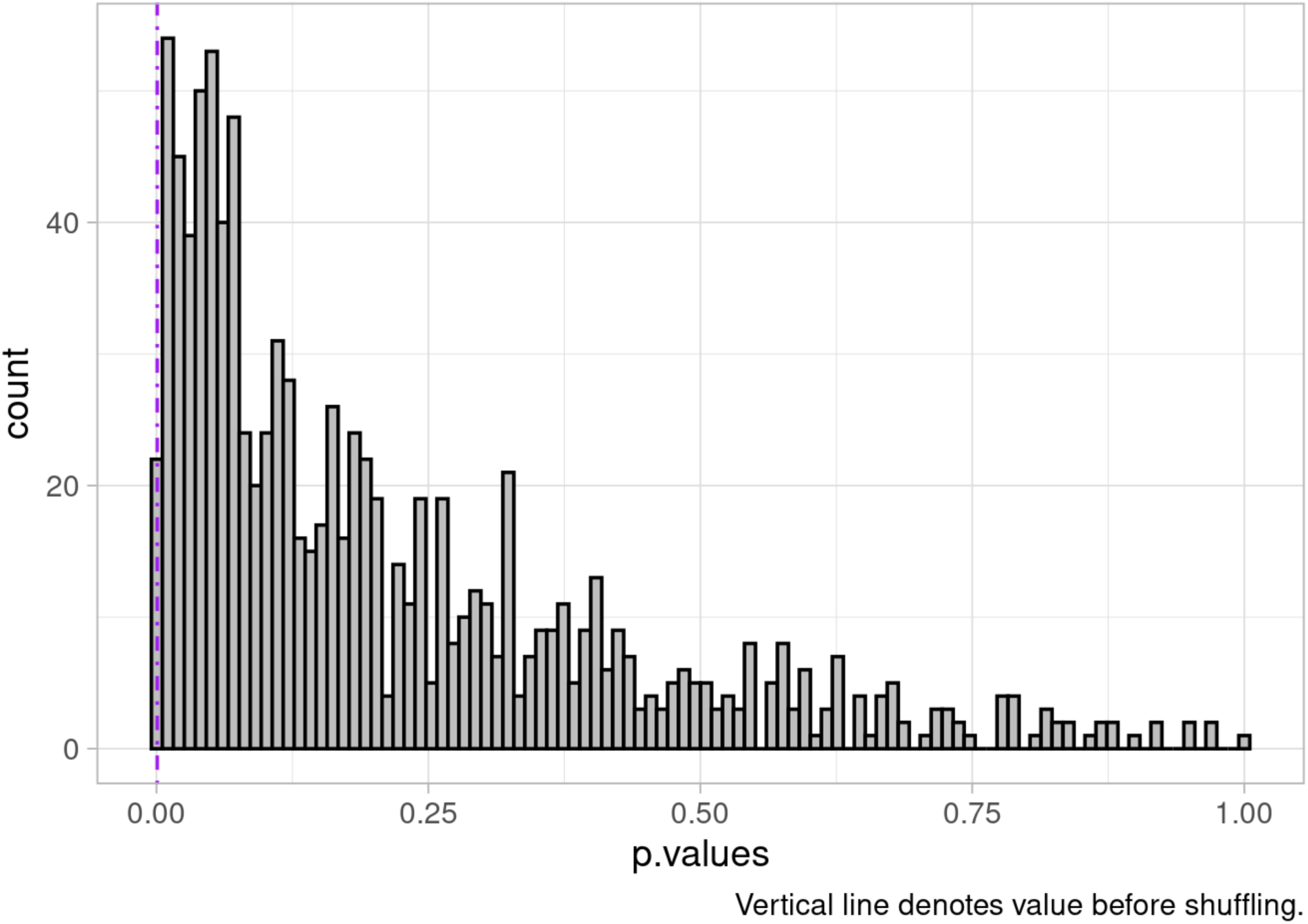
Null distribution of p-values based on 1000 iterations batch-label permutation followed by ComBat batch correction and Wilcoxon rank sum test. Only one iteration produced a p-value as significant as the outcome of the batch correction applied to the study cohort indicating the reliance of batch correction on our experimental batches for correction resulting in strong discriminative ability. This outcome provides confidence that batch correction behaves as expected and does not spuriously introduce differences between experimental response classes.

**Figure 7B:**
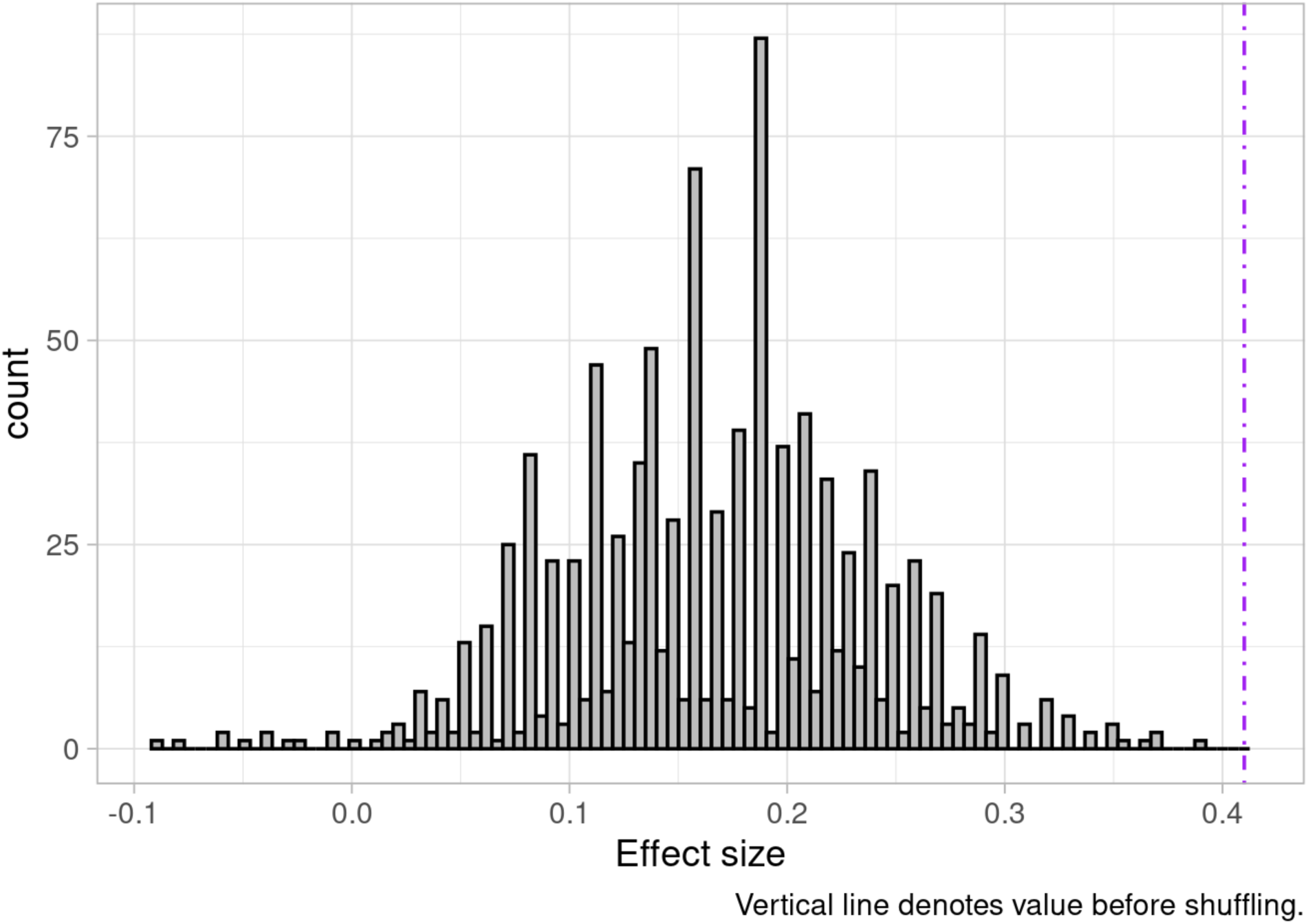
Null distribution of effect sizes based on 1000 iterations batch-label permutation followed by ComBat batch correction. No iteration produced an effect size equal to the outcome of the batch correction applied to the study cohort, indicating the reliance of batch correction on our experimental batches for correction resulting in strong discriminative ability. This outcome provides confidence that batch correction behaves as expected and does not spuriously introduce differences between experimental response classes.

### Disease free survival analysis

Clinical DFS information was available for most patients. While our cohort was modestly sized and relatively underpowered for a time-to-event analysis we generated an exploratory analysis using Kaplan Meier plots to assess the potential for patient-tumor derived MOS to predict longer-term patient outcomes. This analysis was not stratified by study phase and was only conducted for the full study cohort due to the limitations of sample numbers and the high potential for spurious results in underpowered data subsets. Post-batch compensation results (Figure 8) showed convincing separation (log rank test p=0.18) and higher MOS assay responsiveness appeared to correctly predict longer DFS. While the analysis was exploratory, underpowered for statistical significance, and requires follow-up with an expanded patient cohort, the initial results were an encouraging indication of the potential for our assay to predict time-to-event as well as tumor-level response, and they also provide an orthogonal indication of the ability of batch compensation to correctly attenuate experimental batch effects.

**Figure 8:**
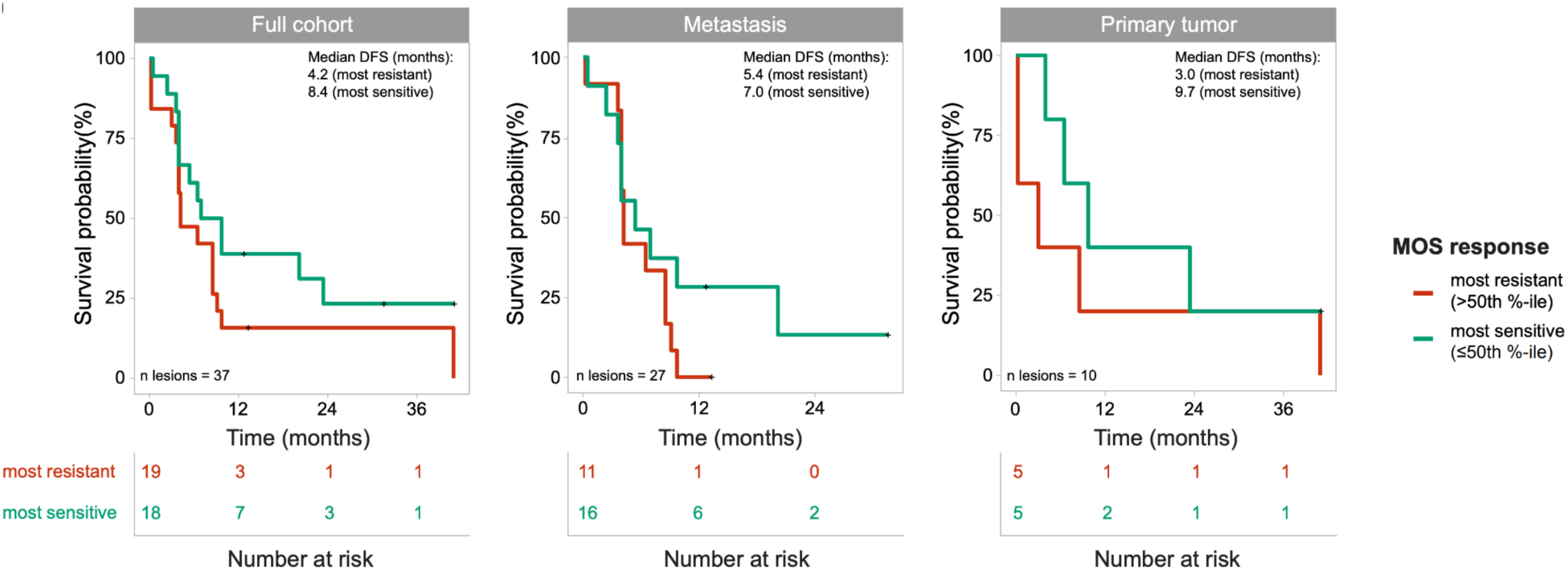
Kaplan-Meier curves depicting the DFS in the full cohort (n = 21 patients), metastasis (n = 16 patients), and primary tumor (n = 9 patients) groups, using a cutoff at the 50th percentile of ranked pIC50s. Groups were compared using the Mantel-Cox log-rank test, and *P* values were adjusted for multiple comparisons using the Benjamini-Hochberg method. **Reproduced with permission from Wolters Kluwer Health, Inc. From: Gobits R, Schleußner N, Oliver GR, et al. *Functional Precision Medicine Using MicroOrganoSpheres for Treatment Response Prediction in Advanced Colorectal Cancer*. JCO Precision Oncology. 2026;10(10):e2500501. © American Society of Clinical Oncology.** https://ascopubs.org/doi/10.1200/PO-25-00501 **This figure is reproduced for non-commercial use in this preprint. No data values or analytical content have been modified. This figure is not covered by the Creative Commons license applied to this preprint.**

Pre-batch compensated data (Figure 9) showed weak separation (log rank test p=0.73) of the patients with the most responsive (>= 50th percentile responsiveness) and least responsive (< 50th percentile of responsiveness) MOS assay results. Furthermore, the visual separation of the data was weakly suggestive of less responsive MOS assays having higher time to event, which is the opposite of what a predictive assay would be expected to indicate. This provided further indication of the strength, necessity and technical grounding of batch-compensation as a first step in the analytical pipeline.

**Figure 9:**
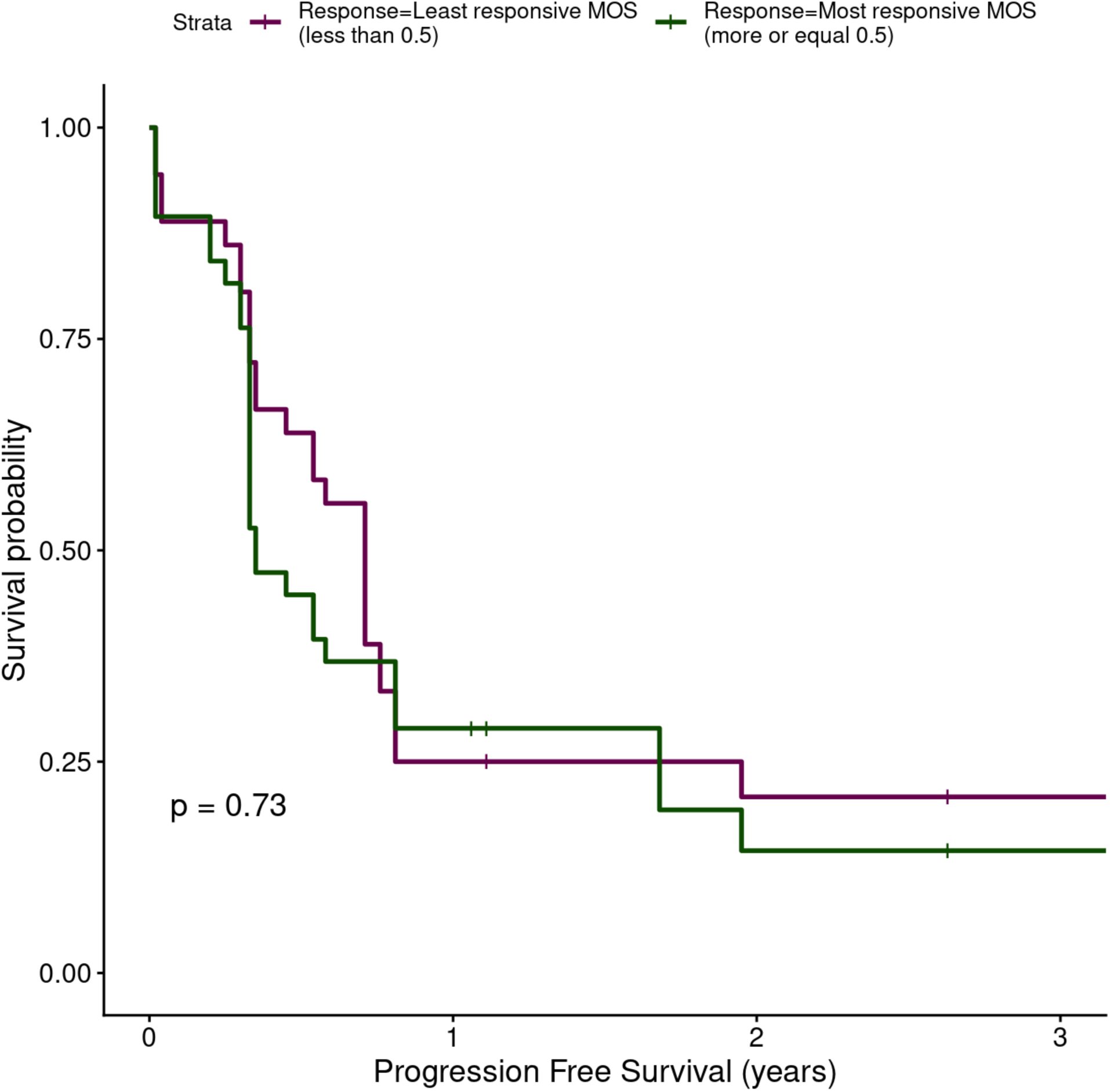
Kaplan Meier plot for all samples (n=37) using where lines represent patients with the most responsive (>= 50th percentile responsiveness) and least responsive (< 50th percentile of responsiveness) MOS assay pIC50s pre-batch compensation. Separation is degraded without batch compensation (log rank test p=0.73) and less responsive MOS appear to show marginally increased survival probability based on visual inspection alone, in opposition to what would be expected if the assay were discriminative.

## Discussion

We have described an approach to monitoring, detecting and compensating for batch effects in a real-world organoid-based retrospective clinical correlation study. By conducting longitudinal monitoring of drug sensitivities using a moving average ^42–47^ to compensate for natural observed variability across our project timeline, and by investigating clinical correlation at the post and pre-batch compensation stage, we were able to identify drift within an initially discriminative assay. Elements of good study design including metadata collection and semi-random order of sample processing subsequently enabled us to investigate and compensate for the issues observed. Application of a traditional method of batch compensation ^48^ with documented wide-ranging use cases ^49–53^, parameterized with recorded batch metadata was capable of attenuating observed batch effects and improving discriminative ability in latter study phases, which matched the observations of the initial study phase, prior to manifestation of overt batch effects. The post-batch compensation results were subjected to extended permutation analysis that demonstrated the robustness of the results and reinforced the basis of batch compensation outcomes in the compensatory importance of the experimental batches themselves, and the improbability of undesired behavior in the batch correction methodology ^58^.

It is clear from the observations we have described that even a carefully conducted study can experience batch-related issues, and diligence is required at all stages of the study, through initial design, execution and analysis. Many good experimental practices as described in our manuscript and in greater detail elsewhere ^15,19^ were implemented and paramount in compensating for batch effects in our study. Nonetheless, occasional shortcomings in design and implementation present learning opportunities for the future. Examples include the fact that media batch information was not recorded in the initial phase of our study and that drug batch information went uncaptured. A further opportunity for future design robustification will be the comprehensive, study-wide use of technical control organoids, or bridging samples ^59^. These were only introduced in the latter phases of our study and could not cover the entire repertoire of agents being tested simultaneously, therefore they provided an incomplete longitudinal control. The information provided by a complete set of control lines would undoubtedly be of high value ^60,61^ and we encourage others to run these alongside study samples with at least one fully representative complement of drugs and doses per experimental batch.

It is likely relevant that our study, which was performed in a recently established laboratory, unavoidably encountered elements of protocol adjustments, equipment and supply chain events that were naturally in flux, increasing the potential for variability to occur even in the presence of operator diligence. Despite trained staff, a high degree of automation in organoid generation, handling, imaging and drug treatment, and widespread use of standard operating procedures, batch effects nonetheless manifested and demonstrated the capacity to compromise the signal of interest across two of our study phases. Notably, within an operations environment we have processed and treated control organoid lines across many months without notable drift in drug sensitivity being detected (data not shown), demonstrating the robustness of the technology when combined with fully established protocols.

As stated, the issues observed in our study occurred despite good practices, state-of-the art technology, and automation being in-place. Inevitably others engaged in organoid studies will face increased challenges introduced by operating within less controlled, well-equipped or automated environments. Here, the potential for issues to arise will only increase. Furthermore, as industry and academia migrate increasingly toward the use of NAMs ^62^, organoids will grow in ubiquity, and the number of studies, publications and confounded results will inevitably expand alongside them. It is vital that investigators recognize the potential for confounded results and adapt suitable best practices within their studies ^13,14,19,55,57^.

The lack of consideration given to batch control in organoid studies is notable in the face of their recognized potential for variability ^4,11,28–30,32^ and this challenge is evidenced by the effects we have reported in a single study of modest scale. The literature appears effectively devoid of studies dedicated to characterizing batch effects in organoid studies, or organoid studies that at least methodologically detail if and how they consider the potential for batch effects. As noted by others ^57^ there appears to be a tendency to condense description of batch effect treatment in published works to the point of uninformativeness, and we believe it is necessary for mandatory minimum documentation frameworks to be formulated and adopted, particularly in light of increasing clinical or clinically adjacent use-cases. The discussion of batches and their treatment within these studies should be brought to the fore as key requirements of study documentation and academic publications. Ultimately recognizing these issues and bringing them into the open will facilitate collaborative efforts, improve understanding and by extension lead to dedicated studies and peer-reviewed or consensus-led optimized methods and protocols for their avoidance or correction. At the very least, describing the methods used to process study data is a long-recognized best practice and if conducted, will provide others with the ability to investigate, repeat and potentially refine such approaches ^63^.

Regardless of best intentions and adherence to good practices, it is well recognized that a real-world clinical study will present challenges that may be difficult to entirely overcome ^19,64^. Even in collaboration with a clinical center of excellence, the order and number in which samples are obtained can be unpredictable. This can lead to the potential for imbalance in study design, unintended temporal clustering of experimental groups of interest, or inconsistencies in batch sizes. Collaboration with external partners also has the potential to introduce a disconnect between clinical and research teams. With limitations in control acknowledged, it is important to ensure that cross-team conversations and planning commence early and continue regularly. The potential for experimental confounding to occur should be discussed and efforts should be made to ensure that all sites are following standardized protocols, recognized metadata points are known and recorded, and that all measures possible are taken to ensure some extent of randomization in the order of sample group processing, and the formation of batches of an acceptable size. Changes to protocols or deviations from standard practice should be communicated and noted centrally alongside sample metadata to ensure its availability at analysis time. Superfluous information can ultimately be filtered or ignored, but missing data can rarely be restored.

Within the course of our study, it was not possible to identify with certainty the cause of the observed batch effects. While batch metadata in combination with appropriate software was capable of attenuating the effects, the batch itself is likely only correlative with the underlying cause. Our observed effects point toward a longitudinal increase or decrease in drug potency or sensitivity depending on the phase and drug, but whether this was caused by differences in the treatments themselves or another factor remains undetermined. While drug batch was not recorded, standard practice dictated that drugs be discarded following three freeze-thaw cycles and follow up diagnostic experiments were unable to produce reduced drug potency in control line organoids following up to ten cycles (data not shown). The inability to fully diagnose a cause versus identify a surrogate in instances of batch correction is a known and accepted limitation ^19^ of scientific experimentation, since it is generally not possible to consider or record every variable of potential interest. As an example, a study of microarray assays was able to show that something as theoretically benign as atmospheric ozone levels in fact had an insidious ability to affect experimental variability ^65^. This goes to further enforce the need to plan and capture information as widely as possible throughout a study timeline, so that causes or surrogates are captured as completely as possible and follow-up investigations and analyses are empowered. While there is always room for improvement, our record keeping was sufficiently comprehensive to enable retrospective investigation of our data, exploration of univariate linear trends, assessment of a set of possible confounders, and ultimately the parameterization of batch compensation software with the processing batch IDs that enabled attenuation of the batch effects.

We have intentionally described our experience in the form of a narrative, rather than attempting to frame it as a protocol. Others’ studies will likely possess their own nuances, and require study-specific customization of approaches to deal with batch effects. Furthermore the underlying cause of an issue may be unique enough to necessitate customized analysis or corrective action. Our approach is not intended to be exhaustive but we believe it represents an initial blueprint that others might study and optimize for their purposes. We hope that the description of our approach will both raise awareness and bestow others with a solid foundation to base their analytical frameworks upon, particularly when such a dearth of literature exists on the topic. We believe that this discussion can provide a valuable and overdue consideration of the topic and seed further literature discussion that will lead to novel and robust methodologies and packages specific to organoid studies in a similar manner to which these have been developed for other areas of science.

## Methods

### Dose response analysis

Sample quality control and dose response analysis were conducted as described previously ^39^. In brief, MOS droplets were individually segmented in brightfield microscopy images and EpCAM fluorescence signal quantified per-MOS droplet using computational analysis. Endpoint signal was normalized by the initial timepoint signal to normalize for variations in biomass. Whole-well signal was calculated as the sum of individual MOS droplet signals per-well. Dose response curves were constructed using a 4-parameter log logistic fit with the DRDA R package ^66^. The curve was normalized to the median vehicle and kill control well signals. Curves were manually inspected for quality control. Notably assays were treated with multiple drugs in order to provide increased data points for assay monitoring and batch-compensation. Samples treated clinically with FOLFOX also had FOLIRI or FOLFOXIRI included in their assay (50/50 split approximately). Samples treated clinically with FOLFIRI received both FOLFIRI and FOLFOX in their assay. Finally, samples treated clinically with FOLFOXIRI received FOLFOXIRI and FOLFOX in their assay, with one case also receiving FOLFIRI.

### Rolling mean analysis

Rolling-mean analysis was performed using the rollapply function from the R zoo package ^67^. A window size of 3 was selected to compute right-aligned rolling means, enabling the inclusion of single observations at the beginning of each data series. Single observation inclusion was achieved using the partial = TRUE argument, which enables windows with fewer than three observations to be evaluated.

### Linear regression analysis

Sequential univariate linear regression models were generated, stratified by drug and phase. Separate models were fit for each candidate confounder. Models were fit using the lm function in the base R stats package. Each model regressed pIC50 on a single independent variable. Coefficient estimate, confidence interval, and p-value were extracted and ranked by p-value to identify the confounders associated most strongly with pIC50.

### Tumor-level clinical correlation analysis

Clinical correlation was performed as described previously ^39^ but in brief, a percentile-based MOS drug response score was calculated for each treatment category using pIC50 values (pre and post batch compensation). Scores were combined across treatments for inter-group comparison. Binary tumor-level clinical response was determined by combining radiological response for metastatic tumors (≥20% growth = Progression, consistent with RECIST thresholds) with pathological Dworak score (TRG0 = Progression; TRG1–4 = Response). Wilcoxon rank-sum tests were used to compare response groups, and p-values were adjusted for multiple comparisons using the Benjamini–Hochberg method. To account for non-independence of multiple lesions contributed by the same patient, we two-sided p-values from 2,000 patient-level permutations in which responder status labels were randomly reassigned to patients (and their lesions) under the null-hypothesis of no association. The final p-value corresponded to the proportion of permuted statistics as or more extreme than the observed value.

### Receiver operating curve analysis

Receiver operating characteristic (ROC) analysis was performed using the pROC package ^68^ in R to evaluate the ability of pIC50 values to discriminate clinical response. Area under the curve (AUC) was calculated with 95% confidence intervals estimated by bootstrap resampling under a binormal model. Uncertainty around sensitivity and specificity were calculated using a nonparametric, patient-level cluster bootstrap. Sensitivity, specificity, and accuracy were recalculated at a fixed classification threshold for each bootstrap dataset and 95% confidence intervals calculated for 2000 replicates. The classification threshold was derived from ROC analysis where candidate thresholds were ranked by balanced accuracy (average of sensitivity and specificity). In cases where the threshold with the highest accuracy corresponded to extreme imbalance (e.g. near-perfect sensitivity but poor specificity), the next highest accuracy threshold that provided a more balanced trade-off between sensitivity and specificity was selected.

### Batch correction

Batch compensation for pIC50 values was performed for each drug individually using the ComBat algorithm from the sva package in R ^48^. The processing batch was provided as the batch label (`batch = processing_batch`). ComBat was run in mean-only mode (`mean.only=TRUE`) and responder status was supplied as a biological covariate (`mod = model.matrix(∼responder_status, data = pIC50)`).

e.g.

~~~
ComBat(dat = data_matrix, batch = batch_factor,mod = model.matrix(∼responder_status, data = pIC50),mean.only = TRUE)
~~~

### Permutation analysis

Batch labels were randomly shuffled among samples before the full batch correction process was rerun using the newly labeled samples and the Wilcoxon rank sum test was applied using the wilcox.test function in base R, and effect size calculated. This analysis was repeated for 1000 iterations, generating empirical null distributions, and the location of the original batch correction p-value and effect sizes in the null distribution were calculated.

### Time-to-event analysis

Individual MOS drug response scores were correlated with patient-level disease-free survival (DFS) using Kaplan–Meier visualizations to explore trends in time to progression. A 50th percentile cutoff of the MOS Response Score was used to split the cohort. The log rank test was run to determine significance of group differences. Survival curves and log-rank p-values were generated using the survminer::ggsurvplot function ^69^ in R.

